# Beaked and Killer Whales Show How Collective Prey Behaviour Foils Acoustic Predators

**DOI:** 10.1101/303743

**Authors:** Natacha Aguilar de Soto, Fleur Visser, Peter Madsen, Peter Tyack, Graeme Ruxton, Patricia Arranz, Jesús Alcazar, Mark Johnson

**Affiliations:** BIOECOMAC. Dept. Animal Biology, Edaphology and Geology. University of La Laguna. Tenerife. Canary Islands. Spain.; Scottish Marine Institute. University of St. Andrews. Scotland. UK.; University of Liege. Netherlands.; Dept. Ecophysiology. University of Aarhus. Denmark; Sea Mammal Research Unit. University of St. Andrews. Scotland. UK.

## Abstract

Animals aggregate to obtain a range of fitness benefits, but a common cost of aggregation is increased detection by predators. Here we show that, in contrast to visual and chemical signallers, aggregated acoustic signallers need not face higher predator encounter rate. This is the case for prey groups that synchronize vocal behaviour but have negligible signal time-overlap in their vocalizations. Beaked whales tagged with sound and movement loggers exemplify this scenario: they precisely synchronize group vocal and diving activity but produce non-overlapping short acoustic cues. They combine this with acoustic hiding when within reach of eavesdropping predators to effectively annul the cost of aggregation for predation risk from their main predator, the killer whale. We generalize this finding in a mathematical model that predicts the key parameters that social vocal prey, which are widespread across taxa and ecosystems, can use to mitigate detection by eavesdropping predators.

## INTRODUCTION

Vital functions such as courtship and foraging are mediated by acoustic signals in taxa as diverse as humans and insects^1^. However, sound-signallers must trade off the benefits of detection by intended receivers against the costs of detection by eavesdropping predators. Strategies for reconciling these conflicting selection pressures remain largely unexplored for sound signals in stark contrast to the intensive study of visual ecology^2^. A common strategy of many prey is to aggregate to reduce risk of predation via dilution or confusion effects^3,4^. These benefits are partially offset by the cost of larger aggregations being more detectable to predators from a distance^3,5^, but the maximum detection distance typically rises sub-linearly with group size. In chemically-or visually-mediated systems the relation between group size and maximum detection range scales with a power between 0.5 and 1^6,7^, but a general relationship for the scaling factor for acoustic cues has not been established. This is surprising given that collective acoustic signalling is widespread in nature and chorusing has been observed in many invertebrates, fish, amphibians, birds and mammals, in both terrestrial and aquatic environments^1,8^.

The intuitive expectation that a larger number of vocal prey will unavoidably enlarge the acoustic detection range of a group may not always be true. In the case of chemical cues, increasing group size enlarges detection distance because the higher concentration of chemicals means that detection thresholds will be met at larger convective distances^6^. Similarly, enlarged visual cues arising from prey aggregation increase maximum detection ranges^8,9^. In contrast, the acoustic source level of aggregated vocal animals only increases if their sound cues overlap in time, similarly to intermittent and short duty cycle (proportion of time that the signal is on) visual cues, such as the flashes of non-synchronized fireflies^10^. Aggregated vocal individuals that are vulnerable to predation should adopt strategies that maximise their cumulative effect on legitimate receivers^11^ but minimise reception by eavesdropping predators. Defining these strategies and how they depend on the characteristics of the habitat and the functions of vocal signals is essential to understand sound-mediated prey predator interactions that are ubiquitous in nature.

Toothed whales provide an ideal case-study to investigate acoustic predator-prey interactions given their reliance on active acoustic detection (echolocation) and passive listening to hunt and sample their environment^12^. Predation pressure from acoustic-guided killer whales (*Orcinus orca*)^13^ has been proposed as an evolutionary driver for the vocal behaviour of the multiple small toothed whale species that produce cryptic high frequency calls, out of the main spectral band of sensitivity of killer whales: Phocoenidae, Kogiidae, and species of genus *Cephalorhynchus*^14^. In contrast, larger species forming tight social groups such as female-young sperm whales (*Physeter microcephalus*)^15^ and pilot whales (*Globicephala* spp)^16^ seem to rely on social defences to abate killer whale predation risk^17,18^. This strategy is not practical for medium-sized beaked whales (Ziphiidae)^13^ which form small social groups and suffer killer whale predation in a wide latitudinal range^13,19^. This source of mortality can be critical for slow-reproducing beaked whales and thus may constitute a strong evolutive force on the behaviour of these deep-diving species.

As in myriad other social animals, aggregation dilutes individual predation risk to beaked whales. Killer whales, the main predator of beaked whales, seem to require the combined efforts of several individuals to subdue a single whale prey^13,19^, providing opportunities for other beaked whales in the group to escape. But the net benefit of aggregation would reduce if aggregated beaked whales are more detectable by killer whales. Here we use novel biologging data from beaked whales to study how their social behaviour affects encounter probability with killer whales. Beaked whales feed using echolocation signals^20^ that can be heard by killer whales. They forage alone or in groups and only vocalise when deeper than 200-500m in deep dives^21^. At these depths they are safe from predation because the short dives of killer whales are insufficient to subdue a beaked whale at depth. However, beaked whales are vulnerable to attack when they surface to breathe if killer whales can locate and track them through a dive. Here we show that a finely-tuned combination of collective behaviours and acoustic hiding by beaked whales reduces by >90% their encounter probability with killer whales, regardless of beaked whale group size. In comparison, continuous and uncoordinated group vocalization would lead to near-certain post-detection interception of beaked whales by killer whales. We generalise these results to model the general principles of abatement of acoustically mediated predation risk by any vocal prey (Box 1), showing that vocal animals can benefit from aggregation while avoiding the penalty of increased acoustic detectability in larger groups.

## RESULTS

### The killer whale-beaked whale acoustic predator-prey system

In predator-prey systems, the temporal and spatial availability of prey cues are key factors influencing detection rate of prey by predators. Here, vocal and diving behaviour data from 27 Cuvier´s and Blainville´s beaked whales obtained with suction-cup attached sound and movement recording tags (DTAGs^22^) (SI) are used to investigate how group size influences beaked whale cue rate and spatial footprint and thus detection probability by killer whales.

Beaked whales are coined extreme divers because they perform stereotyped diving cycles day and night comprising a deep and long foraging dive with maximum duration and length of 2 hrs and 3 km (Cuvier´s beaked whale), followed by a series of shorter and shallower recovery dives separated by brief (mode∼2.5min) surface intervals to breath^23–25^. Individual beaked whales are vocal on average 18%-20% of their time, for echolocation and occasional social signalling during deep foraging dives ^21,26^. Beaked whales are typically found at the surface in tight groups although these groups lack long-term stability. We tagged pairs of whales in the same social group in three instances finding remarkable activity synchronization within these three whale pairs (Figure 1 and SI Table 1). While animals were within a group, the most coordinated deep dives (defined as the two deep dives with closest start time performed by the two whales in each whale pair) overlapped on average for 99% of dive duration (SD 0.3%). The vocalisation phase of such dives overlapped in time by 98% (SD 4%). The most coordinated shallow dives overlapped by a mean of 97% (SD 2.4%). A randomization test showed that in 100% of 4000 iterations the observed dive-profiles rendered a higher overlap of dives than simulated data obtained by random permutation of the dive cycles of one of the whales of the pair (SI). Real overlap exceeded random overlap by an average of 44% (SD 24%) of the time in both deep and shallow dives, and by 63% (SD 31%) of the vocal phase time (SI).

**Fig. 1:**
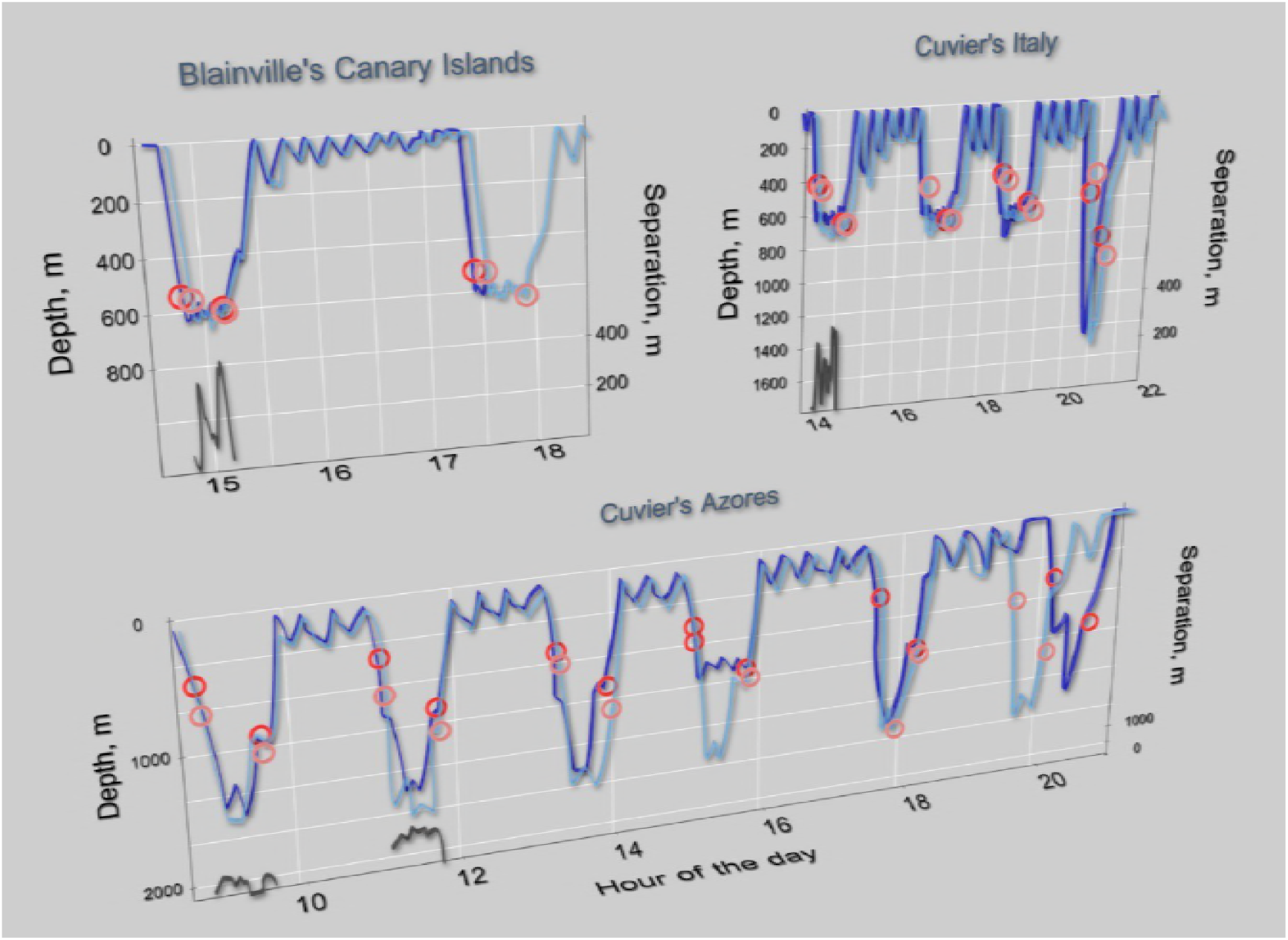
Dive profiles of three pairs of whales tagged in the same social group, showing in light and dark blue the dives of each whale of each pair. A) Two Blainville´s beaked whales in the Canary Islands; B) and C) Two Cuvier´s beaked whales tagged in Italy and Azores, respectively. The group in the Azores was observed to split after the 5th deep dive and there is no further diving coordination after the split. The circles mark the start and end of the vocal phase of each animal in the dives. The black lines at the base of the dives indicate the separation distance between animals in a pair during the vocal phase of these dives.

**Supplementary Table 1.** Dive coordination of the three pairs of whales tagged simultaneously in the same social group. Md: Blainville´s beaked whales, *Mesoplodon densirostris* tagged off El Hierro, Canary Islands; Zc Genoa: Cuvier´s beaked whales, *Ziphius cavirostris*, tagged in the Ligurian Sea (Italy); Zc Azores: Cuvier´s beaked whales tagged off the Azores. Information is pooled for the dive pairs (i.e., the two dives with closest start time performed by the two whales of the pair) performed by each whale pair: Max. depth and Depth diff.: mean of the maximum depth of the two dives of each dive pair and difference in the maximum depths of the dives within each dive pair (m). Dur. and Dur. diff.: mean duration of, and mean difference in, the duration of the two dives of each dive pair (min). Time overlap: mean of the proportion of time that the two dives in dive pairs overlap with respect to the duration of each one of these dives. Vocal overlap: mean of the proportion of time that the vocal phase of the two dives in dive pairs overlap with respect to the duration of the vocal phase of each one of these dives. All data are expressed as mean (range) pooling the results of all dive pairs for each whale pair.

| Whale pair | Dive pairs | Max. depth (m) | Depth diff (m) | Dur (min) | Dur diff (min) | Time overlap (%) | Vocal overlap (%) |
| --- | --- | --- | --- | --- | --- | --- | --- |
| Md Hierro | Deep n=1 | 639 | 16 | 46 | 2 | 99 | 98 |
|  | Shallow n=6 | 56 (37-108) | 8 (2-13) | 11 (9-14) | 0.7 (0.5-1.2) | 93 (91-96) | - |
| Zc Geneva | Deep n=4 | 954 (724-1600) | 49 (6-135) | 55 (48-64) | 0.2 (0.2) | 100 (99-100) | 95 (90-100) |
|  | Shallow n=11 | 173 (114-275) | 21 (6-42) | 17 (15-20) | 0.3 (0.1-1) | 98 (94-100) | - |
| Zc Azores (all dives) | Deep n=6 | 1568 (1296-1670) | 229 (89-769) | 61 (58-62) | 0.9 (0.1-4) | 88 (29-100) | 83 (0-100) |
|  | Shallow n=15 | 162 (142-189) | 39 (22-61) | 19 (16-22) | 0.6 (0.2-0.8) | 97 (95-99) | - |
| Zc Azores (pre-group split) | Deep n=5 | 1544 (1296-1669) | 287 (89-769) | 61 (60-62) | 0.3 (0.1-0.6) | 100 (100-100) | 100 (100-100) |
|  | Shallow n=12 | 166 (142-189) | 39 (22-61) | 19 (16-22) | 0.6 (0.2-0.8) | 97 (95-98) | - |

Similar group vocal coordination was observed in an additional dataset of 54 deep vocal dives from 12 whales tagged separately in different groups. The mean duration of the vocal phase in these dives was 25 minutes. The time-delay of start/end of clicking between the tagged whale and any conspecific whale within acoustic range of the tag differed by just 1.8 min (SD 1.5, start of clicking) and 0.9 min (SD 1, end of clicking) (Supp. Table 2). These results for single tagged whales in groups from 2 to 6 whales are consistent with the observed 98% overlap in the vocal phase of dives performed by paired tagged whales (SI Table 1).

**Supplementary Table 2.** Difference in the timing of start and end of clicking (SOC and EOC, respectively) between tagged Blainville’s beaked whales and any untagged whale within acoustic range of the tags. Results are provided in minutes and expressed as the mean (std) for each tag deployment. The name of the tag deployment is codified with the two last digits of the year, the Julian day of the deployment and a letter indicating the consecutive tag order of the day. In some cases, clicks from other animals could not be assessed due to elevated background noise (primarily flow noise on tags located posteriorly in the whale) or EOC could not be assessed because the tag released before the end of the dive; in these cases the number of dives used for analysis is reported in brackets.

| Whale | # vocal dives | Duration vocal phase | Time-diff SOC | Time-diff EOC |
| --- | --- | --- | --- | --- |
| Md03_284a | 6 | 26.23 (4.9) | 2.31 (1.21) | 0.75 (1.34) |
| Md03_298a | 2 | 24.79 (3.07) | 0.05 (0.06) | 0.35 (0.14) |
| Md04_287a | 4 | 27.51 (4.22) | 0.65 (0.8) | 0.23 (0.21) |
| Md05_277a | 3 | 25.38 (3.24) | 2.03 (0.31) | 1.06 (0.59) |
| Md05_285a | 4 | 25.11 (2.18) | 2.5 (1.63) | 0.99 (1.17) |
| Md05_294a | 2 (1) | 21.04 | 0.43 (0.38) | 0.09 |
| Md05_294b | 4 | 20.87 (2.4) | 1.87 (1.71) | 0.67 (0.36) |
| Md08_136a | 2 | 24.32 (3.33) | 0.73 (0.32) | 0.39 (0.16) |
| Md08_137a | 8 | 27.95 (5.87) | 5.9 (4.74) | 1.42 (0.78) |
| Md08_142a | 2 (1) | 20.42 | 1.82 (0.66) | 0.26 |
| Md08_148a | 2 (1) | 27.18 (7.74) | 1.53 | 4 (4.5) |
| Md08_289a | 7 | 26.18 (9.11) | 1.82 (1.33) | 0.73 (0.49) |
| Md10_146a | 1 | 21.85 | 1.48 | 0.81 |
| Md10_163a | 7 | 20.5 (4.67) | 0.75 (0.79) | 0.2 (0.18) |

Adding the mean observed offset in clicking timing of group members to the mean duration of the vocal phase of tagged whales results in a mean of 27.7 min of group vocal activity per dive. Thus, considering the mean dive cycle duration of 120-140 min^23,26^, groups of whales are acoustically available for detection some 20-22% of their time. This is only slightly longer than the 18-20% of time that individual whales within a group are available for acoustic detection^21,26^, meaning that the proportion of time that beaked whales are available for passive acoustic detection by killer whales is almost independent of group size. In comparison, a randomization test simulating a signalling channel with activity slots that can be accessed by one or more whales at random predicts an approximately Gaussian distribution for the time that 6 asynchronous beaked whales would be available for acoustic detection. The mean of this distribution is 69%, i.e. more than three times longer than the observed 22% of the time that a group of six beaked whales is vocally active, showing how vocal coordination can reduce the time that animals are available for predator detection.

Animals with highly synchronized vocal activity will reduce the time availability of their acoustic cues to potential predators, but this may happen at the cost of increasing spatial availability. This depends on the vocal duty cycle of the animals, i.e. the proportion of time that animals are signalling within a vocal period, and thus the probability of signal overlap. The detection range of acoustic cues increases when cues overlap in time and their power sums, (e.g. chorusing frogs^1^). However, the probability of vocal cue overlap in beaked whales is extremely low even when individuals in social groups synchronize the vocal phase of their dives. This is because apart from rare short whistles^21^, beaked whales only produce short (∼200 μs) echolocation clicks with a mean duty cycle of 0.0007^21^. Moreover, the volume of water ensonified by the highly directional clicks of beaked whales^27–28^ increases negligibly in groups. This is because beaked whales diving in tight coordination show a similar circular distribution of the pointing angle of their clicks within a dive, (i.e. they ensonify a similar restricted sector of the circle) (SI).

Inter-animal separation also influences cue spatial availability. Groups cannot be considered an acoustic point source when they disperse. We calculated the separation between pairs of beaked whales tagged simultaneously in the same group using an acoustic travel-time method (SI). Whales were as close as 11 m when they began echolocating at a mean depth of 450 m. They then separated by up to 1500 m while hunting but re-joined at the end of the vocal phase to as close as 28 m before initiating the silent ascent from a mean depth of 760 m (Figure 1). Taken together, the whale pairs spent 95% of the vocal phase less than 500 m apart. Considering an individual on-axis maximum detection range of 6.5km^29,30^, and the typical 90º coverage of clicks within a dive, the separation of 0.5 km between beaked whales in a group means an increase in the detection area for surface-dwelling killer whales of 16% of a group compared to a single beaked whale.

In sum, the collective diving and vocal behaviour of beaked whales reduces cue time availability by 40% and increases detection footprint by just 16% while still allowing animals to disperse to hunt. This increase in spatial detectability given by group dispersal occurs when beaked whales are at depths that provide them a refuge from shallow diving killer whales. However, diving beaked whales are susceptible to acoustic stalking in which killer whales track them acoustically and then attack when they leave their deep-water refuge during obligate surfacing for air. Here, the collective behaviour of beaked whales is key to foil stalking predators. By coordinating their dives, groups of diving beaked whales are released from a “surface anchor” that would be maintained by the need to re-join with non-diving group members and thus frees groups to choose where to surface from dives. Most deep-diving whales ascend steeply to minimize transport time and hence maximize foraging time at depth^31,32^, however, this behaviour leads to a high encounter probability with killer whales stalking acoustically from the surface. In contrast, both Cuvier´s and Blainville´s beaked whales manoeuvre in a way that confounds surface predators when they ascend to breathe. These whales silence at an average depth of 760 m and ascend towards the surface with an unpredictable heading and a shallow average pitch angle of 35º with respect to the horizontal^23,33^. This unusual behaviour for an air-breathing mammal creates an uncertainty cone for the position of beaked whales while they ascend in silence. The resulting potential surfacing area is a circle of 3.7 km^2^ (∼1.1 km radius) centred on the position of the last click emitted by diving beaked whales (Fig 2 and SI).

**Figure 2:**
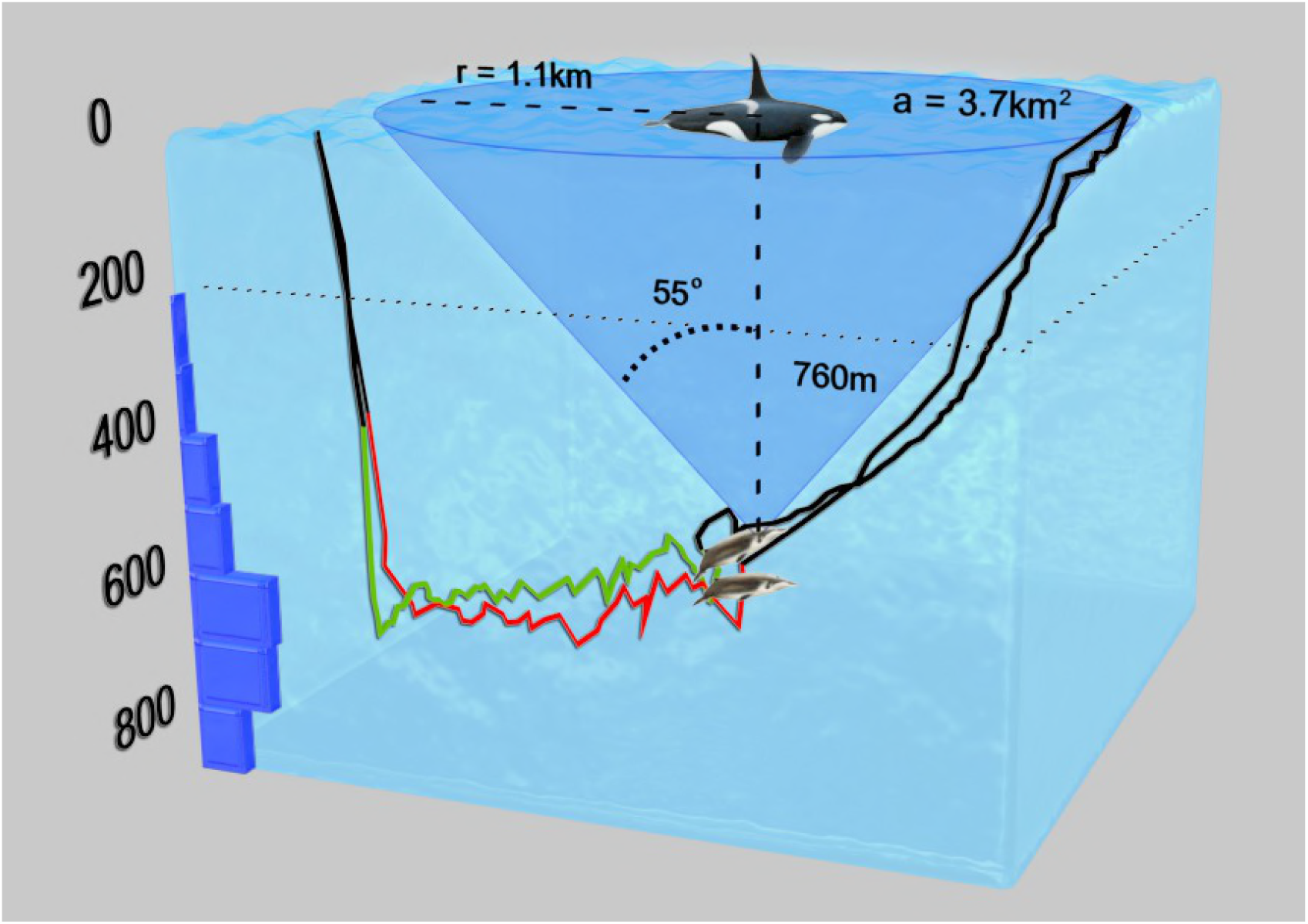
Post-detection encounter probability is <10% for killer whales acoustically stalking beaked whales due to the uncertainty in their surfacing location following long silent ascents. The coloured lines in the dive profiles of two beaked whales diving in coordination represent the vocal phase of these dives. The histogram is the depth distribution of the clicks of beaked whales (truncated to 900 m), showing that they are silent at the depths to which killer whales usually dive (marked as a dotted line at 200 m depth).

A pod of killer whales that has tracked acoustically deep diving beaked whales could potentially dive to hunt the beaked whales at depth. However, this does not seem feasible given the protracted and intense pack hunting effort required for killer whales to subdue cetaceans at the surface^13,19^, and the restricted 10 min duration of killer whale dives^34^. Thus, killer whales need to wait for beaked whales to be at or near the surface to hunt them. Killer whales are unlikely to use echolocation to track beaked whales to avoid alerting them and elicit avoidance responses^35,36^. This means that killer whales must search visually the uncertainty surfacing area of beaked whales in the short time that beaked whales spend at the surface after a vocal dive, before they dive again. Both Cuvier´s and Blainville´s beaked whales spend a median of 2.5 min at the surface after a dive and this short surfacing is typically followed by a relatively shallow and tightly coordinated silent dive of up to 400 m depth and 25 min duration^23^ in which beaked whales can again move hundreds of metres horizontally. Assuming a usual swimming speed of killer whales of 2 m/s^37^ and a visual detection range of some 50 m underwater, an individual killer whale can cover visually only some 0.6% of the potential surfacing area of beaked whales during the 2.5 min that beaked whales are at the surface. Encounter probability increases with killer whale pack size: usual pack size of mammal eating killer whales is 3-4 whales, but up to 12 whales have been observed^19^. Killer whales in large packs and perfectly coordinated to not overlap in search area could cover some 7% of the potential surfacing area of beaked whales.

Thus, the coordinated movement and acoustic hiding behaviour of Cuvier´s and Blainville´s beaked whales results in a maximum probability of interception by stalking predators of 7% irrespective of group size, i.e., a reduction of >90% when compared to the high interception probability for animals that ascend vertically and/or vocalise during the ascent. The unpredictable ascent of beaked whales is only possible due to their coordinated diving behaviour.

## DISCUSSION

Beaked whales exemplify a widespread strategy of vocal animals: to broadcast when predators are not detected or when in a safe place with limited predator access (e.g. in the case of beaked whales, deep waters are safe from killer whale attacks), and silence (i.e. hide acoustically) when compelled to leave the refuge or when predators are detected. These behaviours are observed in avian nestlings, as well as in chorusing insects and frogs, that silence in response to alarm calls or predator approaches^39,40^. Another important commonality among beaked whales and other vocal species is that long-range broadcasting is necessary to achieve the biological functions of echolocation and many communication signals^1^. For all vocal prey, there is a clear evolutionary bonus in reducing predation risk while fulfilling these biological functions.

The results of this paper show that the detectability of beaked whales for their main natural predator, the killer whale, is very similar for individuals and groups. Tagged beaked whales emitted on average 41% (∼1500 clicks) of the clicks produced in a dive while the whales were oriented towards the sea surface, at an average rate of 68 (SD 22) upward clicks per min of the vocal phase. This means that killer whales crossing the acoustic footprint of beaked whales at slow speeds of less than 2 m/s^38^ have a high probability of detecting a single vocalising beaked whale when passing by the ensonified area, and thus additional clicks from several vocal whales with collective vocal behaviour may be redundant for group location. In contrast, vocal group size will likely influence beaked whale detection probability from non-natural receivers passing at faster speeds, such as ships with hydrophone systems. Natural predators such as killer have limited capacity to swim faster for protracted times to increase their search area, but they would improve their encounter rate of beaked whales by increasing group size and spreading out while performing area restricted search of detected beaked whales. In fact, killer whale groups attacking beaked whales are larger than groups attacking other marine mammals^19^, indicating that cooperative searching is one way that killer whales can combat the abatement tactics of beaked whales.

In addition to predator defence, coordinated diving may provide additional benefits to beaked whales. An advantage could be sharing information^41^ via eavesdropping on the foraging activity of group members as has been observed in echolocating bats^42^. Coarse level local enhancement is important when groups forage in patchy resources and beaked whales may be attracted to richer patches indicated by the acoustically determined prey encounter rate of their group members. However, we show here that beaked whales do not appear to forage cooperatively regularly, because individuals disperse several hundreds of metres during the echolocation phase of the dive. Simultaneous diving in absence of coordinated foraging has been observed in other air-breathing vertebrates, such as penguins^43^, where this collective behaviour provides a further example of the benefit of aggregation to dilute predation risk.

The extraordinary collective behaviour of beaked whales and its clear benefits for predation risk abatement led us to generalise the results by constructing a quantitative model of the parameters influencing acoustic predation risk abatement. The opportunities and strategies available for vocal animals to abate acoustically mediated predation risk depend on the functions and characteristics of their vocalizations, the acoustic transmission properties of the medium, and the movement patterns and group behaviours associated with sound production. In Box 1 we present a general model that demonstrates how vocal group size affects predation risk for any vocal animal in terrestrial and marine environments.

The model in Box 1 illustrates that low duty cycle animals that call asynchronously such as echolocators strongly reduce their predation risk in terms of reduced detectability by aggregating. In contrast, aggregated animals vocalizing with high time overlap (whether because of a high duty cycle or precise synchronization) do not reduce detectability when transmitting in environments in which sound spreads spherically such that signals decrease in intensity with the inverse of distance-squared. Further, they incur an enhanced detectability when vocalising in conditions of cylindrical spreading (i.e., in which signals decrease with the inverse of distance, such as in shallow water or temperature inversions^44^). These cases of geometrical spreading and animal vocal synchronicity frame a range of potential intermediate scenarios in nature. Thus, the model summarises the main parameters influencing the strategies available to abate acoustically mediated predation risk for any gregarious vocal prey. These parameters are activity synchronization, vocal time-overlap, group aggregation and habitat sound transmitting properties.

We have presented scenarios encompassing a range of potential outcomes of animal behaviour and habitat characteristics on the active acoustic space of vocal fauna. In an extreme (but not far-fetched) case we predict that there is little difference between the acoustic detectability of a single individual and of a tight group of animals with synchronous vocal periods but no overlap in vocalizations; the killer whale-beaked whale predator-prey system exemplifies this strategy. In contrast, detection range is amplified by increasing time-overlap of calls and vocal group cohesion in habitats where geometric spreading loss tends towards cylindrical models. Increased predation risk may be a necessary cost of the fitness advantages provided by long-range vocal signalling, but observation of inheritable behavioural tactics reducing predation risk in obligate sound producers^45^ underlines the importance of reducing the risk of detection in the evolution of animal vocal behaviour.

## BOX 1

### General principles of prey behaviour for abatement of acoustic predation risk

In acoustic predator-prey interactions, prey detection by predators is a probabilistic function of the proportion of time in which acoustic cues of prey are available to predators (T), and of the spatial footprint of these cues (S). Animal groups can reduce T by synchronizing individual periods of vocal activity. This tactic, observed here in beaked whales, is also exemplified by choruses. An additional benefit of this strategy is the possibility to concentrate vocal activity to periods in which predators are absent or prey are in locations safer from predators. A cost of synchronising general vocal activity for predation risk is a higher probability of time overlap of individual calls increasing S, as is the case in choruses^1,3^. Thus, animals may trade the anti-predator benefits of a reduced T for the predation costs of an increased S. Moreover, animals may use vocal synchronization intentionally to extend S, e.g. chorusing in periods when climatic conditions such as thermal inversion favour reception by intended receivers^1^. Surprisingly, a larger S may not linearly increase predation risk in some cases, e.g. frog-eating bats respond less to synchronous than asynchronous frog calls^46^. This might be explained by the confusion effect of simultaneous signalling frogs making it difficult for bats to resolve the angle of arrival of individual calls and locate the emitter. In these cases, prey benefit from reducing the time they are available for detection by predators, without paying the full cost of an increased detection footprint.

The effect of vocal group size on S varies for different animals and habitats. Here we derive a simplified general model applicable to any vocal species to investigate the effect of vocal group size on acoustic detectability. For a group of *n* vocal individuals, we term *n_s_* as the number of individuals with synchronized, time-overlapping, vocal cues. The model is derived for two vocal strategies: asynchrony of vocalizations of individual group members (i.e., stochastic channel access), and full time-overlap of individual vocalizations. Denoting individual duty cycle as *d*, the vocal strategy modulates *n_s_* as follows:

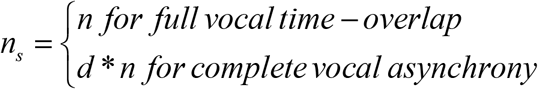

The effect of increased *n_s_* on *S* depends on the acoustic transmission loss (TL) in the broadcasting habitat and on the geometry of the detection footprint. TL is dominated by geometric spreading loss and other attenuation effects of sound energy, such as absorption and scattering^1^. Absorption is most relevant at high frequencies^44^ although in terrestrial habitats vegetation acts as a band pass filter^47^. Because absorption and other sound attenuation effects, but not geometric spreading loss, are frequency dependant^1,44,47^, here we construct a simple model applicable to all signals and habitats, to investigate the relative effect of group size on detectability under different types of geometric spreading transmission loss (TL) and summarise the effects of absorption as a multiplicative (additive in Decibels) term *a* (SI). Geometric TL fits or is intermediate between cylindrical and spherical models in most habitats, i.e., TL (Decibels) ∼ G*log_10_(r)+a, where G equals 10 and 20 for cylindrical and spherical loss, respectively^1,44^. A general relation between the maximum detection range of a group, r_group_, and an individual, r_ind_, is the following (derivation in SI):

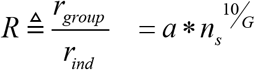

Modelling the widespread and simplified case of a circular detection area results in the following relations among R, S and *n_s_* for different sound transmitting habitats within the extremes of spherical and cylindrical spreading loss (SI). Here, R and S are the ratio of group maximum detection range and acoustic footprint, respectively, with respect to the values of these parameters for an individual:

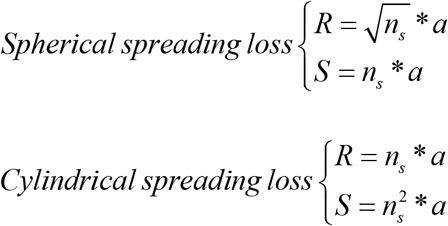

From the above we see that S depends on 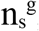, where g=1 in spherical transmission loss and g=2 in cylindrical transmission loss, with intermediate values for other types of geometric spreading loss.

An additional parameter influencing *S* is the dispersion of vocal animals. Tight groups, where the separation among animals is negligible with respect to their individual detection range, function as an acoustic point source. As individuals disperse they enlarge the active space of the group, to the extreme that the acoustic space of a dispersed group with no overlap in the acoustic space of its members is the sum of the acoustic space of all vocal group members. Here we define parameters s_ind_d as the acoustic footprint of an individual; S_ga_ is the acoustic footprint of a group of closely aggregated animals; and S_gd_ is the acoustic footprint of dispersed animals. The combined effects of aggregation and of vocal duty cycle (which influences the probability of signal overlap and thus SL) determine S. This in turn defines the benefit of aggregation for predation risk abatement, defined as B=S_gd_/S_ga_, for groups of animals with different group size, vocal strategies and vocalising in different habitats, as follows:

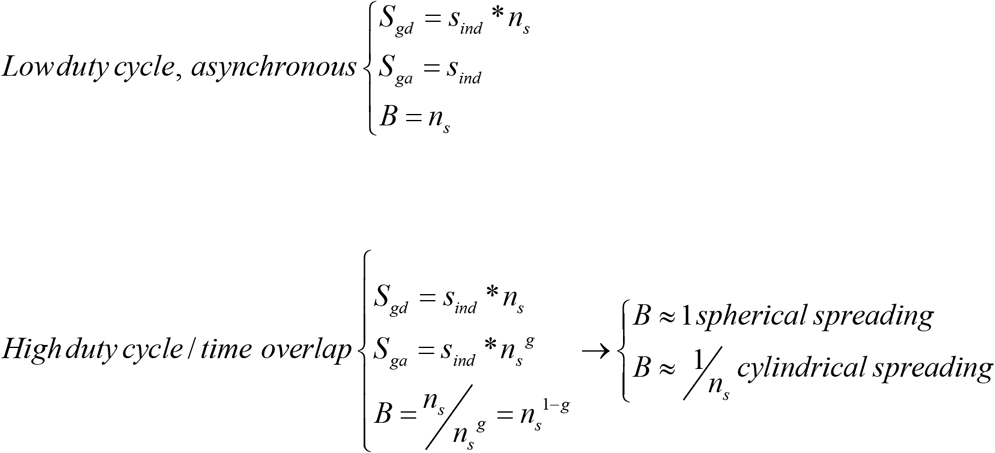

## Acknowledgements

Thanks to all researchers and students collaborating in data collection. This work was funded by the Office of Naval Research (ONR) with support from the Spanish Ministries MAGRAMA and MICINM and a Marie Curie Sklodowska fellowship (EU Horizon 2020, project ECOSOUND). NAS was funded by the later and a by Ramon y Cajal fellowship (MICINM)

## Author contributions

NAS, MJ, PM, FV, PA performed the experiments, JA contributed to the data analysis. NAS and MJ wrote the manuscript with contributions from all authors.

## Supplementary experimental procedures

### Data collection

Beaked whales were studied using suction-cup attached DTAGs^16^ containing depth and orientation sensors (3-axis accelerometers and magnetometers) sampled at 50 or 200 Hz and two hydrophones sampled at 96, 192, or 240 kHz. Blainville´s beaked whales (*Mesoplodon densirostris*, n=14), were tagged off El Hierro (Canary Islands, Spain, see^15^); Cuvier´s beaked whales (*Ziphius cavirostris*) were tagged in the Gulf of Genoa (Ligurian Sea, Italy, see^17^), n=10, and off Terceira (Azores, Portugal, with similar SI as used in El Hierro), n=2. In all cases whales were approached slowly from a small boat and the tag was deployed on the back of the whales with the aid of a handheld pole. Tags were located for recovery using VHF tracking after their programmed release from the whales.

### Tag data analysis

Tag data were analysed in Matlab (*Mathworks*). Depth and whale movement data were calibrated with standard procedures^16^. Sound recordings were examined with custom tools from the DTAG toolbox (www.soundtags.org) to identify vocalizations of the whales. Vocalizations comprised echolocation clicks and buzzes^22^, as well as rasps and rarely whistles with an apparent communication function^15^. Echolocation clicks were located individually with the aid of a supervised click detector^22^.

Cuvier´s and Blainville´s beaked whales perform deep and long foraging dives (deeper than 500m^17^) interspersed with series of shallow dives defined as dives between 20 and 500 m depth^17^. Surfacing intervals separating consecutive dives (both deep and shallow) were measured in the depth profiles. Results were analysed per individual and then averaged for each species. When two whales were tagged simultaneously in the same group (see below), we only used data from the first tag deployment of the pair. Surface intervals lasted on average 2.5 min (std 0.6) and 2.6 min (std 1.3) for Cuvier´s and Blainville´s beaked whales, respectively (mean of the median duration of the surface intervals performed by each whale, grouped by species).

#### Diving and vocal coordination

Groups of beaked whales were defined as clusters of whales observed together at the surface. No inferences were made about short or long-term group stability. Whales in these clusters were most often observed to surface together for the duration of the visual follow. In three occasions (one per field site) we tagged two whales in the same social group. Tag deployments on the two members of each of these three whale-pairs overlapped in time during 3, 9 and 12 hours, respectively; the 6 whales forming these whale-pairs performed in total 22 deep and 64 shallow dives SI (Table 1).

Dive coordination of the whales in whale-pairs was assessed by comparing timing and depth of the most coordinated dives performed by the two members of each whale-pair. These coordinated dive-pairs were defined as the dives with closest start time performed by the two whales of each whale-pair. The analysis was performed separately for deep vocal dives (deeper than 500 m maximum depth) and shallower non-echolocating dives^17^. For the resulting dive-pairs we calculated the time overlap of the dives, as well as the overlap in the vocal phase of vocal (deep) dives. Differences in duration and maximum depth between the dives in each dive-pair were recorded also. Results were pooled for each whale-pair (SI Table 1) and then for the three whale-pairs given the close similarity in results between study areas and species and the small sample size of Blainville´s beaked whales (all but one dive-pairs were recorded from Cuvier´s beaked whales).

The group of Cuvier´s beaked whales tagged in the Azores was followed by the research boat and observed at a distance during surfacing intervals to monitor group composition via individual photo-identification. Analysis of photographic data showed that the four animals forming the group at the time of tagging continued to surface in close vicinity until some 9.5 hrs after tag deployment. After this, two of the four whales, including one of the tagged whales, were no longer observed in the group. The analysis of dive coordination of this Azorean whale-pair was performed both for the full duration of the double tag deployment and for the time before the group split (SI Table 1).

A randomization test was performed to estimate the likelihood of the observed overlap of dive-pairs occurring by chance. For each whale-pair we compared the overlap in observed dive-pairs, against the overlap in simulated dive profiles. Simulations were constructed for each whale-pair using the recorded dive profile of the first tagged whale, and randomly permutated dives from the dive profile of the second tagged whale. The analysis was performed separately for deep and shallow dives, and for the vocal phase of deep dives, with 1000 randomizations for each case. For deep dives, the permutation unit was a deep dive cycle comprising a deep dive and the following interdive interval, i.e. the period of shallow diving before the next deep dive^6,7^. For shallow dives, the permutation unit comprised a shallow dive and its following inter-dive interval (i.e., until the next dive, shallow or deep). The randomization test was not applied to the pair of Blainville´s beaked whales because these whales only shared one full deep-dive cycle, nor to the time after the Azorean group split.

The separation distance between whales in each whale-pair was estimated during the vocal phase of tagged whales. This was achieved by measuring the time delay between the emission of a click by a tagged whale and the reception of the same click on the tag carried by the other whale in the pair. Comparison of time delays for clicks produced by each of the two whales allowed for estimation of the clock offset between the two tags. Clock offset was subtracted from the measured time delays to give the acoustic time of flight which was then converted to distance by multiplying by the path-integrated sound speed, using custom scripts from the dtag-toolbox (www.soundtags.org, M. Johnson). Depth profiles of sound speed for each location were used together with the known depths of each animal to derive the path-integrated sound speed for each click. Sound speed profiles were gathered from CTD (RBR Ltd. and Sea-bird Scientific Inc.) deployments performed at El Hierro and the Ligurian Sea at the time of tagging, and from the AZODC database for Azores (http://oceano.horta.uac.pt/azodc/oceatlas.php) in a relatively close area and season of the year with respect to the tagging event.

### Paired tagged whales click directionality

Because echolocation clicks are highly directional, group size could increase detectability if whales ensonify their surroundings at random. We performed a circular analysis of the heading of the whales while producing clicks shows that whales in a group tend to ensonify a very similar circular sector within each dive (SI Fig. 1)

**Figure SI:**
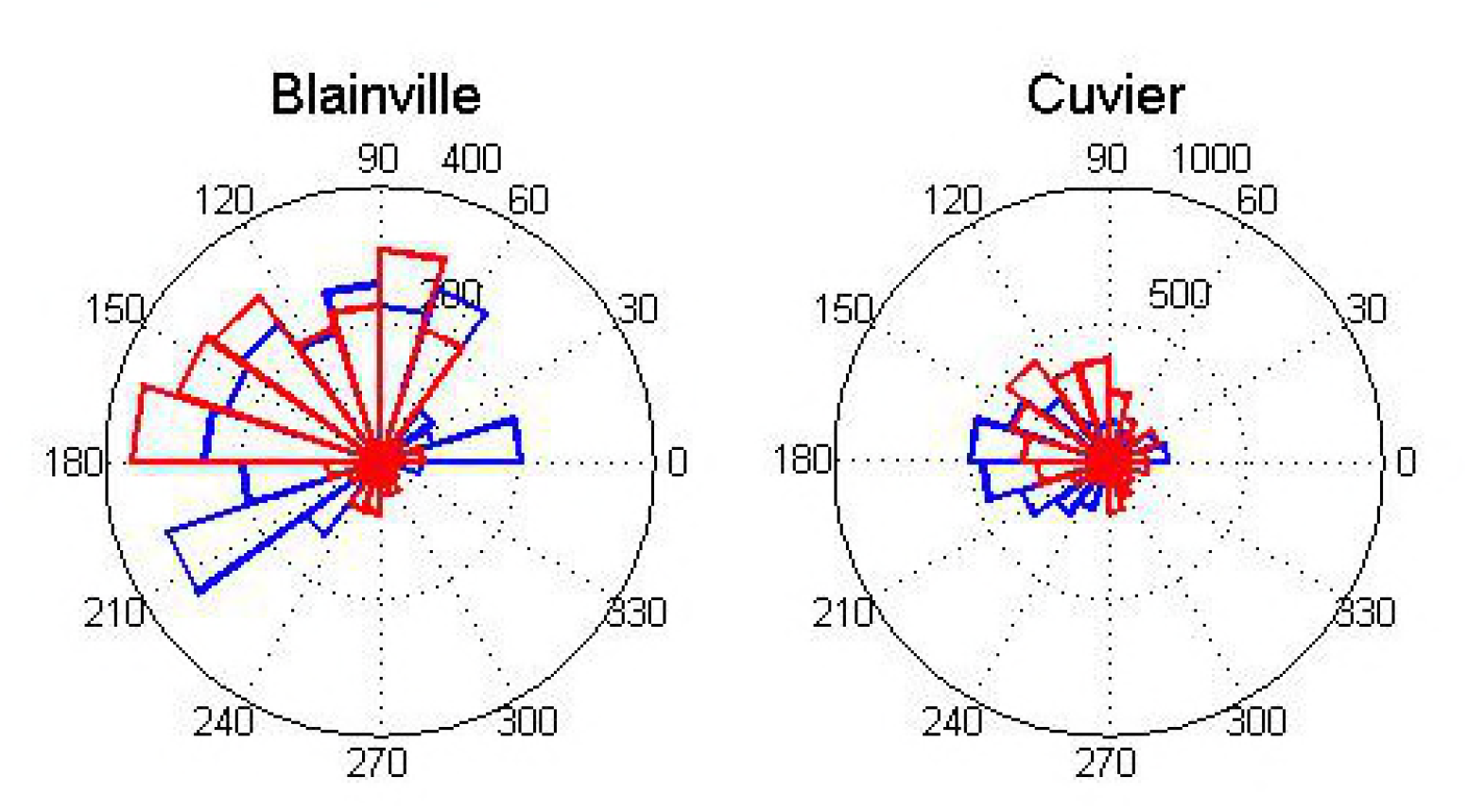
Example of the circular distribution of the heading of the whales while producing clicks in one dive. Each rose shows the results for a pair of whales tagged in the same social group (in red and blue for the two members of the whale pair) performing near-simultaneous dives.

### Calculation of search surface area for killer whales

Tagged beaked whales ended clicking on average at 760 m depth and ascended with a mean pitch angle of 35º with respect to the horizontal^27^, i.e. 55º with respect to the vertical. This renders a maximum surfacing area described by the base of a cone with height equal to the depth of the whale at the time of silencing and a half internal angle of α= 55º. This potential surfacing circle has a radius r=1085 m (r=h*tan(α)) and an area a=3.7 km^2^ (a=π*r^2^). These are maximum values if whales maintain a constant heading during the dive ascent. Previous analysis^17^ have shown that Cuvier´s and Blainville´s beaked whales adopt a fairly constant heading during ascents, covering consistently more than 50% of the maximum horizontal distance assuming a constant heading, and more than 80% of the maximum distance in 55% of the dives^17^. It is possible that beaked whales modulate the horizontal distance covered during ascents according to the distribution of foraging resources and to the presence of predators or other potential disturbing stimuli, such as ships^38^ or delphinids, which have been observed to harass beaked whales (Ana Cañadas, pers.com).

### General acoustic model formulae derivation

In all acoustic detectors, a requisite for detection is that the signal to noise ratio, i.e. the source level (SL) minus the noise level in the area (NL) minus the transmission loss (TL), equals or exceeds a given required detection threshold (DT):

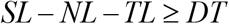

TL can be simplified as the sum of geometrical spreading with coefficient G^35^ and absorption, considering an absorption coefficient α and a maximum detection range r, as follows:

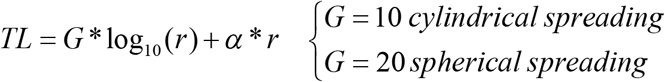

The SL of a group of ns vocally overlapping individuals relates to individual SL as:

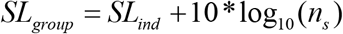

Because the DT required by a predator to detect prey does not depend on prey group size we can solve DT for an individual and for a group and equal them as follows:

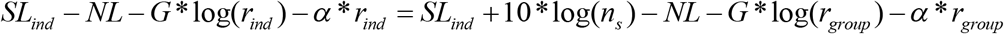

For a given SL_ind_ and NL we can simplify the equation above by dividing by G all elements and expressing them as logarithms to solve the relation *R* between maximum detection range for a group and an individual, as follows:

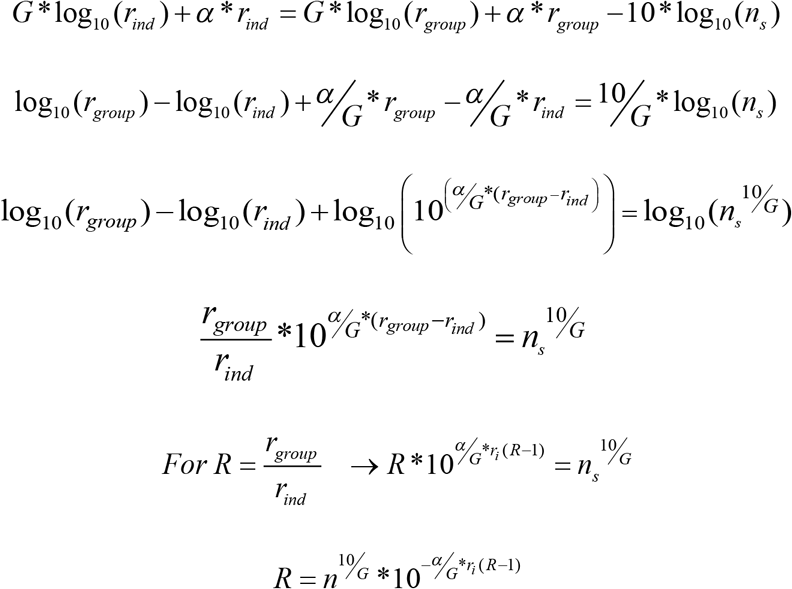

We will term the effects of absorption *a*, so that: 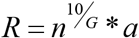

In many cases the receiver is constrained to a 2-dimensional search surface (e.g., shallow water predators eavesdropping on a deep-water caller, or terrestrial animals searching for prey on the ground) and this renders a circular detection area. This results in the following relations between the maximum range (R) and area (S) of detection of a group of *n_s_* overlapping vocal animals with respect to an individual, for different sound transmitting habitats within the extremes of spherical and cylindrical spreading loss:

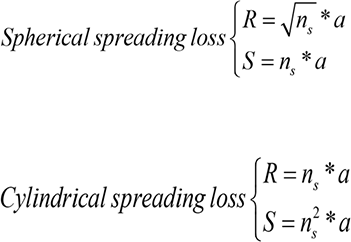

**Supplementary video 1:** Two-dimensional animation of the dive profile of two Blainville´s beaked whales tagged in the same group, in blue and black, showing the start and end of the vocal phase of the dive of each animal with asterisks. The video evidences the high coordination of the diving and vocal behaviour of the whales. The animation runs 40 times faster than the real data.

**Supplementary video 2:** Tagging of beaked whales and animation of their diving behaviour including DTAG data on the vocalizations of the whales. Video courtesy of St. Thomas Productions, part of the documentary “Champions of the deep” (http://www.saint-thomas.net/uk-program-81-marine-mammals-champions-of-the-deep.html).

## References

1. Burt, J. M. & Vehrencamp, S. L. Animal Communication Networks. (Cambridge University press, 2005).

2. Cronin, T. W., Johnsen, S., Marshall, N. J. & Warrant, E. J. Visual ecology. (Princeton University Press, 2014).

3. Krause, J. & Ruxton, G. D. Living in Groups. (Oxford University Press, 2002).

4. Sword, G. A., Lorch, P. D. & Gwynne, D. T. Insect behaviour: migratory bands give crickets protection. Nature 433, 703–703 (2005).

5. Mchich, R., Auger, P. & Lett, C. Effects of aggregative and solitary individual behaviors on the dynamics of predator-prey game models. Ecol. Model. 197, 281–289 (2006).

6. Andersson, P., Löfstedt, C. & Hambäck, P. A. How insects sense olfactory patches - the spatial scaling of olfactory information. Oikos 122, 1009–1016 (2013).

7. Low, C. Grouping increases visual detection risk by specialist parasitoids. Behav. Ecol. 19, 532–538. (2008)

8. Radford, C., Stanley, J., Tindle, C., Montgomery, J. & Jeffs, A. Localised coastal habitats have distinct underwater sound signatures. Mar. Ecol. Prog. Ser. 401, 21–29 (2010).

9. Bukovinszky, T. et al. The role of pre-and post-alighting detection mechanisms in the responses to patch size by specialist herbivores. Oikos 109, 435–446 (2005).

10. Buck, J. & Buck, E. Biology of synchronous flashing of fireflies. Nature 211, 562–564 (1966).

11. Ruxton, G. D., Sherratt, T. N. & Speed, M. P. Avoiding Attack. (Oxford University Press, 2004).

12. Perrin, W. F., Wursig, B. & Thewissen, J. G. M. ‘Hans’. Encyclopedia of Marine Mammals. (Academic Press, 2009).

13. Jefferson, T. A., Stacey, P. J. & Baird, R. W. A review of killer whale interactions with other marine mammals: predation to co-existence. Mammal Rev. 21, 151–180 (1991).

14. Morisaka, T., and R. Connor. Predation by killer whales (*Orcinus orca*) and the evolution of whistle loss and narrow-band high frequency clicks in odontocetes. J. Evol. Bio. 20:1439–1458 (2007)

15. Whitehead, H. Babysitting, dive synchrony, and indications of alloparental care in sperm whales. Behav EcolSociobiol 38, 237–244 (1996). 43.

16. Augusto, J. F., Frasier, T. R. & Whitehead, H. Characterizing alloparental care in the pilot whale (*Globicephala melas*) population that summers off Cape Breton, Nova Scotia, Canada. Mar. Mammal Sci. (2016).

17. Pitman, R. L., Ballance, L. T., Mesnick, S. I. & Chivers, S. J. Killer whale predation on sperm whales: observations and implications. Mar. Mammal Sci. 17, 494–507 (2001).

18. Stephanis, R. D. et al. Mobbing-like behavior by pilot whales towards killer whales: a response to resource competition or perceived predation risk? Acta Etho. 18, 69–78 (2014).

19. Wellard, R. et al. Killer whale (*Orcinus orca*) predation on beaked whales (*Mesoplodon spp.*) in the Bremer Sub-Basin, Western Australia. PLoS ONE 11, e0166670 (2016).

20. Johnson, M., Madsen, P. T., Zimmer, W. M. X., Aguilar de Soto, N. & Tyack, P. L. Beaked whales echolocate on prey. Proc. R. Soc. Lond. BBiol. Sci. 271, S383- S386 (2004).

21. Aguilar de Soto, N., et al. No shallow talk: Cryptic strategy in the vocal communication of Blainville’s beaked whales. Mar. Mammal Sci. 28, E75–E92 (2012).

22. Johnson, M. & Tyack, P. A digital acoustic recording tag for measuring the response of wild marine mammals to sound. IEEE J. Ocean. Eng. 28, 3–12 (2003).

23. Tyack, P. L., Johnson, M., Aguilar de Soto, N., Sturlese, A. & Madsen, P. T. Extreme diving of beaked whales. J. Exp. Biol. 209, 4238–4253 (2006).

24. Baird, R. W. Diving behaviour of Cuvier´s and Blainville´s beaked whales in Hawaï. Can JZool 84, 1120–1128 (2006).

25. Schorr, G. S., Falcone, E. A., Moretti, D. J. & Andrews, R. D. First long-term behavioral records from Cuvier’s beaked whales (*Ziphius cavirostris*) reveal record-breaking dives. PLoS ONE 9, e92633 (2014).

26. Arranz, P. et al. Following a foraging fish-finder: diel habitat use of Blainville’s beaked whales revealed by echolocation. PLoS ONE 6, e28353 (2011).

27. Madsen, P. T. et al. Biosonar performance of foraging beaked whales (*Mesoplodon densirostris*). J. Exp. Biol. 208, 181–194 (2005).

28. Johnson, M. et al. Foraging Blainville’s beaked whales (*Mesoplodon densirostris*) produce distinct click types matched to different phases of echolocation. J. Exp. Biol. 209, 5038–5050 (2006).

29. Zimmer,W. et. al. Echolocation clicks of free-ranging Cuvier’s beaked whales (*Ziphius cavirostris*). J. Acoust. Soc. Am. 117,3919–3927 (2005).

30. Marques, T. A., Thomas, L., Ward, J., DiMarzio, N. & Tyack, P. L. Estimating cetacean population density using fixed passive acoustic sensors: An example with Blainville’s beaked whales. J. Acoust. Soc. Am. 125, 1982–1994 (2009).

31. Miller, P. J. O., Johnson, M. P., Tyack, P. L. & Terray, E. A. Swimming gaits, passive drag and buoyancy of diving sperm whales Physeter macrocephalus. J. Exp. Biol. 207, 1953–1967 (2004).

32. Aguilar Soto, N. et al. Cheetahs of the deep sea: deep foraging sprints in short-finned pilot whales off Tenerife (Canary Islands). J. Anim. Ecol. 77, 936–947 (2008).

33. Martin Lopez, L. M., Miller, P. J. O., Aguilar de Soto, N. & Johnson, M. Gait switches in deep-diving beaked whales: biomechanical strategies for long-duration dives. J. Exp. Biol. 218, 1325–1338 (2015).

34. Miller, P. J. O., Shapiro, A. D. & Deecke, V. B. The diving behaviour of mammal-eating killer whales (*Orcinus orca*): variations with ecological not physiological factors. Can. J. Zool. 88, 1103–1112 (2010).

35. Falcone, E. A. et al. Diving behaviour of Cuvier’s beaked whales exposed to two types of military sonar. R. Soc. Open Sci. 4, 170629 (2017).

36. Barrett-Lennard, L. G., Ford, J. K. B. & Heise, K. A. The mixed blessing of echolocation: differences in sonar use by fish-eating and mammal-eating killer whales. Anim. Behav. 51, 553–565 (1996).

37. Deecke, V. B., Slater, P. J. B. & Ford, J. K. B. Selective habituation shapes acoustic predator recognition in harbour seals. Nature 420, 171–173 (2002).

38. Williams, R. & Noren, D. P. Swimming speed, respiration rate, and estimated cost of transport in adult killer whales. Mar. Mammal Sci. 25, 327–350 (2009).

39. Magrath, R. D., Pitcher, B. J. & Dalziell, A. H. How to be fed but not eaten: nestling responses to parental food calls and the sound of a predator’s footsteps. Anim. Behav. 74, 1117–1129 (2007).

40. Stevens, M. Sensory Ecology, Behaviour, and Evolution. (Oxford University Press, 2013).

41. Danchin, É., Giraldeau, L.-A., Valone, T. J. & Wagner, R. H. Public Information: From Nosy Neighbors to Cultural Evolution. Science 305, 487–491 (2004).

42. Barclay, R. M. R. Interindividual use of echolocation calls: Eavesdropping by bats. Behav. Ecol. Sociobiol. 10, 271–275 (1982).

43. Takahashi, A. etal. Penguin-mounted cameras glimpse underwater group behaviour. Proc. R. Soc. Lond. B Supplement 271, 281–282 (2004).

44. Urick, R. J. Principles of underwater sound. (McGraw-Hill, New York, 1983).

45. Hedrick, A. V. Crickets with extravagant mating songs compensate for predation risk with extra caution. Proc. R. Soc. Lond. B Biol. Sci. 267, 671–675 (2000).

11 Tuttle, M. D. & Ryan, M. J. The role of synchronized calling, ambient light, and ambient noise, in anti-bat-predator behavior of a treefrog. Behav. Ecol. Sociobiol. 125–131 (1982).

46. Aylor, D. Noise reduction by vegetation and ground. J. Acoust. Soc. Am. 51, 197–205 (1972).

